# Negative biodiversity-ecosystem function relationship in broad but not in narrow functions within pitcher plant microbial communities

**DOI:** 10.1101/2025.07.11.664437

**Authors:** Catalina Cuellar-Gempeler, José Alejandro Bravo, Eric Malekos, Dan Perez-Sornia, Marissa Monaco, Sandrine Grandmont-Lemire, Megan Teigen

**Author notes:** **Corresponding author:** Dr. Catalina Cuellar-Gempeler. Address: 1st Harpst Street, Arcata CA. 95521.

## Abstract

The relationship between Biodiversity and Ecosystem Function (BEF) addresses how communities transform their environment. BEF relationships can have positive, neutral, or negative slopes, yet it remains unclear what conditions result in a particular slope. A popular classification in microbial ecology distinguishes ‘broad’ from ‘narrow’ functions and we ask whether this distinction improves predictions of BEF relationships. Specifically, we evaluate whether the relationships between broad functions and diversity can be predicted based on (1) the combined slopes from underlying narrow functions, (2) the phylogenetic breadth of associated species and (3) their ecological dominance. We assembled bacterial communities from pitcher plant fluid, using a dilution-to-extinction approach to create a gradient in biodiversity. *Darlingtonia californica* are carnivorous plants that depend on their bacterial community to degrade insects that supplement their nitrogen requirements. We found a negative BEF relationship between bacterial richness and degradation, while narrow functions had positive and neutral BEF slopes. The narrow functions did not predict the BEF relationship for the broader function. We identified three species statistically associated with degradation: *Clostridium* sp. had a positive association, while *Herbinix* sp. and *Dyadobacter* sp. had negative associations. *Clostridium* sp. was rare in the dataset and negatively correlated with *Herbinix* sp., suggesting an antagonism and highlighting important functional contributions of subordinate species. We propose that the negative BEF for degradation is explained by rare key functional taxa that thrive in low-diversity communities. These findings suggest that species abundance distributions outperform function-function relationships in explaining the emergence of complex broad functions.

**Importance:** In this study, researchers examined how bacterial diversity shapes ecosystem function, using microbial communities from carnivorous pitcher plants as a model system. Contrary to the common belief that higher biodiversity enhances function, they found that communities with higher bacterial richness were *less* effective at breaking down insect prey. While specialized microbial activities improved or remained stable with greater diversity, they did not explain the decline in overall degradation. Instead, a rare *Clostridium* sp. bacterium emerged as a key driver of decomposition in low-diversity communities, while more dominant species were linked to reduced function. These findings reveal that rare microbes can play critical roles in ecosystem processes and suggest that who is present and how abundant they are may matter more than the sum of their potential functions.

## Introduction

Uncovering biodiversity’s role in ecosystem function is vital for conservation and sustainability on a changing planet [1–4]. For decades, studies seemed to support a general positive relationship between biodiversity and ecosystem functions [known as BEF, 5, 6]. Recent publications reveal the commonality of neutral relationships and the rare occurrence of negative BEF relationships [7–9]. An outstanding challenge is to be able to recognize the conditions that lead to positive, neutral, or negative BEF relationships [10, 11].

A potential path forward is to characterize observed functions in a way that forecasts their relationships with diversity and community composition [7, 12–14]. Here, we consider a popular classification system from microbial ecology that distinguishes ‘broad’ from ‘narrow’ functions [15–18]. Broad functions encompass many activities, are performed by multiple community members [15, Table 1], and support large-scale ecosystem processes like respiration, primary productivity, biomass accumulation, and decomposition [12, 19]. Narrow functions are more specific, typically associated with one activity (Table 1), like single carbon substrate utilization [16]. Performed by few community members [15], narrow functions can require large, complex and specific enzymes [12, 19–23] and be limited to narrow phylogenetic group [12, 24, 25], such as cellulose degradation by select cellulase-producing bacteria [26–28].

**Table 1.** Summary of studies comparing broad and narrow functions. Broad and narrow functions are defined for each study according to the system. The BEF relationship is reported for each type of function if it was explicitly tested.

| Broad | Narrow | System | BEF relationship | Reference |
| --- | --- | --- | --- | --- |
| Biomass, C and nutrient flow broadly | Plant microbe interactions (Mycorrhizae, N fixing symbionts), litter decomposition (lignin and cellulose), trace gas production | Tundra, plant microbe interactions, decomposition | Not measured | Schimel 1995 |
| Mineralization of wheat shoot carbon | Mineralization of 2,4 dichlorophenol pollutant | Polluted and agricultural soils | Broad - redundancy, narrow - affected by diversity loss | Girvan et al 2005 |
| Decomposition and organic matter turnover, aerobic heterotrophs | Refers to Schimel 1995 | Soils | Not measured | Schimel and Schaeffer 2012 |
| Respiration | Two toxin degradation functions (x and y) | Fresh water ecosystems | Broad and narrow were positive | Delgado-Baquerizo et al 2016 |
| Mineralization | Nitrification | Degraded soil | Broad and narrow were positive, composition matters | Calderon et al 2017 |
| Respiration, cell yield, ATP concentration | Cleavage rate of 4 substrates (xylose, chitin, cellulose, monoesters) | Beech water filled tree holes | Broad and narrow were positive | Rivett and Bell 2018 |
| Thymidine incorporation, growth decomposition | Nitrification | Soils | Broad - redundancy, narrow - affected by diversity loss | Griffiths et al 2020 |
| Respiration | Degradation of 7 specific substrates | Soils | Broad - redundancy, narrow - variable (positive and neutral) | Li et al 2021 |
| Heterotrophic<br>production and DOC<br>utilization | ECOPLATE-BIOLOG – 31 carbon<br>substrates | Fresh water<br>ecosystems | Broad -<br>redundancy,<br>narrow -<br>affected by<br>diversity loss | Cheng et al 2022 |

Theory suggests that we can predict the general shape of broad and narrow functions. Broad functions are thought to be functionally redundant in diverse communities, such as microbial organisms [29, 30] or tropical rainforests [31], resulting in neutral BEF relationships [32–36]. For instance, many taxa contribute to biomass accumulation or respiration and thus, the loss of diversity should have little impact on the community’s aggregate function [15, 17, 37, 38]. In contrast, narrow functions are expected to decline sharply with biodiversity loss as they rely on a small number of species and are thus more likely than broad functions to have positive BEF relationships [17, 39]. For example, toxin degradation and nitrification were found to have steeper positive relationships with bacterial diversity than the broader functions of respiration and mineralization [24]. An alternative view proposes that narrow functions can have negative relationships with diversity (negative BEF relationships) because they associate with subordinate species that perform poorly in species rich communities (Tranvik and Jorgensen 1995, Jiang et al. 2008, Grover 2009, Peter et al. 2011, Logue et al. 2016).

Contrary to expectations based on theory, broad functions can have positive BEF relationships, while narrow functions can exhibit neutral BEF relationships (Table 1). One explanation is that broad functions act as an umbrella for diverse complementary activities [44–49]. Complementarity effects describes how functionally unique species coexist with minimal overlap in resource use, enhancing functional outputs, likely through facilitation and mutualistic relationships [45, 50]. Other explanations arise from considering aggregate function resulting from redundant, complementary, essential, or antagonistic contributions by narrow functions [51–54]. Additionally, these narrow functions seem to portray a diverse range of BEF relationships, with neutral slopes when functions are shared widely across many groups of taxa [such as the breakdown of common carbohydrates in soils, 17, 38], or negative slopes when they represent energetically costly pathways performed by poor competitors [40]. These diverse patterns indicate that a generalized BEF relationship framework for broad and narrow functions remains unresolved (Table 1).

We address these gaps in our understanding of BEF relationships by evaluating key assumptions underlying the broad and narrow function classification. First, we question the generality of neutral BEF relationships in broad functions, and positive BEF relationships in narrow functions. Second, we ask whether narrow functions can predict broad BEF relationships. Broad functions are sometimes referred to as aggregate functions implying that they emerge from a combination of narrow functions [18, 19]. Third, we establish whether taxa associated with function show ecological dominance, or phylogenetic clustering. Specifically, ecological dominance and broad phylogenetic affiliation have been associated with positive broad BEF functions, while rare, clustered groups are hypothesized to associate with negative narrow BEF relationships.

We investigated the diversity and function of microbial communities associated with the pitcher plant *Darlingtonia californica*. These carnivorous plants attract prey with nectar and trap them in their modified leaves [55, 56]. Because of *D. californica*’s limited enzyme production, bacteria within the pitcher plant fluid play a critical role in the degradation of prey captured by the pitcher shaped leaves [57, 58]. This process of degradation releases nitrogen to the pitcher plan fluid, which is then utilized by the plant to grow in soils with low nutrient concentrations [56, 59]. While nitrogen is a key component for biosynthesis in the plant and the microbes, prey detritus is the only source of carbon for the bacterial community [60]. Therefore, we study the degradation of prey items as the broad function in this system and the subsequent use of carbon as narrow functions that impact the microbial community specifically. Because degradation differs from traditional broad functions like biomass accumulation [7, 61, 62], we anticipate contributing to the landscape of BEF relationships in the context of broad and narrow functions (as in Table 1), while establishing function-function and species-function relationships, enhancing our ability to predict BEF relationships.

## Methods

To evaluate BEF relationships, we created a diversity gradient from pitcher plant fluid using a dilution-to-extinction approach, and measured the degradation function for each of the resulting microbial communities. The dilution-to-extinction is a practical approach to manipulate microbial diversity of environmental samples while keeping complex communities with unculturable taxa [20, 41, 51, 63–66]. We used the 16S rRNA gene to assess bacterial diversity and abundance, and performed a degradation assay to assess function. To establish the generality of our findings, we compare experimental communities with fluid from three populations of *D. californica*.

### Inoculum collection and processing

Pitcher plant fluid was obtained in September 2019, from 3 plants kept in the Cal Poly Humboldt’s Denis K. Walker Greenhouse. We collected fluid and debris from 2 leaves from each plant (6 leaves total) and pooled all contents for a total volume of 12mL. Fluid was transported in ice to the laboratory where it was filtered sequentially through sterile 500um and 350um sieves to remove debris and larger eukaryotic microorganisms. The filtered fluid was brought up to 200mL with sterile DI water and incubated at room temperature with gentle mixing for 8 days (pre-dilution incubation). Field samples were collected in 2021 with similar methods. See supplementary materials for specifics on pre-dilution incubation and field sample collections.

### Experimental set-up

The culture above was used to inoculate three dilutions series by transferring 100mL of the original culture into three sterile 250mL conical tube. Then, we performed five 1:10 dilutions by transferring 10mL of the culture into 90mL of sterile DI water and mixing by inverting 3 times. We created replicates by separating 30mL from each of these dilution tubes into 50mL Falcon tubes, for a total of 45 microcosms. To stabilize the microcosms and standardize bacterial abundances, we incubated at room temperature adding 2mg of sterile ground fruit flies every 2 days. After 4 days, we removed 10 mL to be frozen at −80°C for DNA extractions, 5mL for functional assays, and 1mL to quantify protozoa using contrast microscopy (1mL). The remaining fluid was brought to 15mL with sterile DI water and used for the degradation assay. We included six controls with 30mL sterile DI water, three of which were fed with 3mg sterile ground fruit flies.

### Degradation assay

We conducted an 11-day degradation assay within each of the experimental microcosms. The fruit fly *Drosophila hydei*, common in decaying matter worldwide and in the pet trade, was used as a standardized insect prey to assess the degradation potential of the microbial communities. The animals were obtained from a local pet store, sacrificed by freezing and autoclaved to eliminate their bacterial load. They were then dried for 24 hrs and weighed. They were then placed in a sterile 1.5mL tube with ∼50 holes, small enough to keep the flies in but allow water flow. On average, the beginning weight of the flies was 12 mg within each cage. The cages were then introduced to the microcosm tubes and incubated at room temperature for 11 days, mixing gently daily by inverting the tube twice to increase flow in and out of the caged insect area. At the end of the assay, the fly cage was taken out of each tube with sterile forceps and dried for 24 hrs. The remaining fluid was frozen at −80°C for DNA extractions (10mL). Degradation rate was calculated as weight loss by subtracting fruit fly weight at the beginning of the degradation experiment from fruit fly weight at the end of the degradation experiment.

### Functional assays

We focused narrow functions related to carbon flow as bacteria access the insect hemolymph (simple carbohydrates), nitrogen containing amino acids and amines, phenolic compounds and energy storage (glycogen and lipid)[67–70]. We used Biolog EcoPlates (Biolog Inc. Hayward, CA) to quantify community utilization of 31 independent sources of carbon [71]. This method is limited in its ability to characterize bacterial diversity but can be very powerful to compare functional characteristics and fits the substrates we are looking for [72]. We followed the manufacturer’s protocol and customized as suggested by Leflaive [73, 74]. See supplementary methods for further detail.

### Molecular analyses

Using Qiagen DNeasy PowerWater Kit following the manufacturer’s protocol, we extracted DNA from samples taken from each microcosm before and after the degradation assay. To estimate bacterial diversity, we used 16S rRNA gene amplicon sequencing according to the Earth Microbiome Project protocols [75, 76]. Then, we estimated bacterial abundance by measuring 16S rRNA gene copy numbers using real-time PCR (qPCR) [77]. This method uses ribosomal DNA copy numbers from environmental samples to estimate cell abundance and is widely used in environmental studies [78–80]. Each sample was run in triplicate and results were averaged. See supplementary methods for details on each of the reactions. Sequences can be found in the NCBI Sequence Read Archive under accession number PRJNA1287784.

### Bioinformatics

Raw sequences were demultiplexed, trimmed and quality filtered using standard procedures (see Supplementary materials). ASVs were inferred using the *dada* function in the package ‘dada2’ [version 1.14.1, 81]., with the supporting error model learning (*learnErrors*) and dereplication (*derepFastq*) functions with default settings in the same package. Taxonomy was assigned to the resulting ASVs at the 97% identity using the Silva project genomic database (version 138) as a reference. Based on this ASV table, we calculated the number of species and Pielou’s evenness [82] using functions *specnumber* and *diversity* from the ‘vegan’ package [version 2.6-8, 83].

### Data Analysis

All of these data processing steps and data analysis steps below were conducted in R [84]. All code and metadata are available in the Github page https://github.com/catalicu/DegExp2019.

To determine whether the dilution treatments had an effect on diversity metrics and bacterial abundance, we used a model selection approach informed by the experimental design, and established the best model fit to assess significance. General Linear Models (GLMs) included Gaussian distributions with each metric (richness, or evenness) as dependent factor and dilution factor (log-transformed), dataset (collected pre or post degradation assay) and their interaction as explanatory factors using the base R function glm(). We compared models using Aikaike Information Criterion [AIC henceforth, 85, 86]. The best model was then assessed using the Wald test. The Wald test is a Maximum likelihood approach implemented using the Anova() function in the car package [87]. Based on the results from these models, we focus on richness as a central diversity metric and on communities sampled before the degradation experiment.

We used a similar approach to evaluate BEF relationships between richness and degradation. GLMs with Gaussian distributions included function as response variable and richness as fixed factor. We compared these models to GLMs that included dilution and series as explanatory variables, in addition to diversity, to establish the effect of the dilution treatment and potential variation across series. We used AIC to make comparisons amongst models and the Wald test to assess significant effects. Additionally, to confirm significance, we tested the best performing model against a null model using the function anova() from the car package [version 3.1-3 88]. We repeated this approach to evaluate the relationship between evenness and prey weight loss.

To establish BEF relationship shapes for narrow functions, we repeated this analysis for the 31 carbon sources assay. For these models, we only assessed richness as the diversity metric. Then, we used an Analysis of covariance (ANCOVA) to compare correlations between functions, in an effort to determine which narrow functions may underlie the broad function. Thus, we also compared every function against each other to establish potential functional groups. To illustrate these connections between functions, we used a correlation matrix implementing cor() from the package corrplot [version 0.95, 89]. To establish significance, we used Spearman’s 𝜌 by applying the function cor.mtest().

We used GLMs to search for associations between ASV abundance and each functional metric, as is done for genomic screenings [15, 90]. For this analysis, we selected ASVs with 1000 reads or more across the dataset to avoid missing data issues in the models. This filtering step left us with the 205 most abundant ASVs in the dataset, representing 98.25% of the total abundance in the study. For each ASV, we used a GLM with Gaussian distribution and function as a dependent factor and ASV relative abundance as fixed explanatory factor. We repeated the model to compare across all selected ASVs and functions and performed a Benjamini-Hochberg correction [91, 92] to account for multiple testing. We used a heat map with the function geom_tile() in the ggplot2 package, to illustrate these relationships.

To establish how the overall community composition related to diversity and function, we used multivariate and univariate approaches using functions from the ‘vegan’ package [version 2.6-8 93] and custom scripts. First, we built an NMDS from relative abundance data (k=3) using the metaMDS() function with 100 iterations. We tested for significant differences in community composition based on dilution factor, and series using the function adonis2() from the same package with 999 permutations, and assessing significance for each term. We assessed whether community composition had an impact on degradation using a GLM with Gaussian distribution. We compared models that included the NMDS axis 1 and 2 as well as richness by calculating the AIC and evaluating the best performing model with a Wald test. We also performed a Pearson correlation to confirm the relationship between NMDS axis 1 and weight loss.

Then we established how individual ASV relative abundance responded to the diversity gradient. As above, we selected ASVs with 1000 reads or more to avoid missing data issues. We used GLMs with gaussian distributions that included ASV relative abundance as dependent variable and diversity as fixed factor. To assess significance, we compared models to a null model using the anova() function and adjusted p values for multiple testing using the Benjamini-Hochberg correction.

We compared the diversity and composition in our experiment to field samples. We calculated richness and evenness as above and compared these diversity metrics across datasets using a Kruskal Wallis one-way analysis of variance given the non-parametric distribution of the data. Then, we performed a Dunn test to establish differences between groups.

Finally, to established the effect of dilution treatments on bacterial abundance, measured as 16S rRNA gene copy numbers, we used GLMs with gaussian distributions (see supplementary results).

## Results

Dilutions created a diversity gradient with a range between 16 to 197 ASVs and a mean of 65 ASVs (± 45.445). Dilutions created a weaker gradient in evenness, ranging from 0.268 to 0.766 (mean 0.443±0.113). Because the main diversity gradient was richness, we will focus on this metric for most analyses in this study. Samples taken after the degradation assay were substantially lower in richness (27.5 ± 9.553 ASVs) and evenness (0.443±0.111) and did not show clear diversity gradients (Fig S1), thus we only include them in comparisons with field data. We did not find sufficient protozoa for analyses; thus, these were not included in subsequent analyses (see supplementary results).

### BEF

We found a negative BEF relationship between bacterial richness and the degradation function (Fig 1). The best statistical model to characterize this relationship included ASV richness and series as factors (TableS2a). The Wald test revealed significant effects of richness and series (Table 2) and the model outperformed the null model (F_3_=5.415, p=0.003). Specifically, weight loss declined in 5.85*10^-5^ mg per 10 ASVs added, but this effect varied with series, with series 3 having the strongest slope, followed by series 1 (Fig 1). We did not find a significant relationship between evenness and weight loss (Fig S2) yet, the best performing model outperformed the null model (F_7_=2.424, p=0.039), likely due to the effect of series.

**Figure 1.**
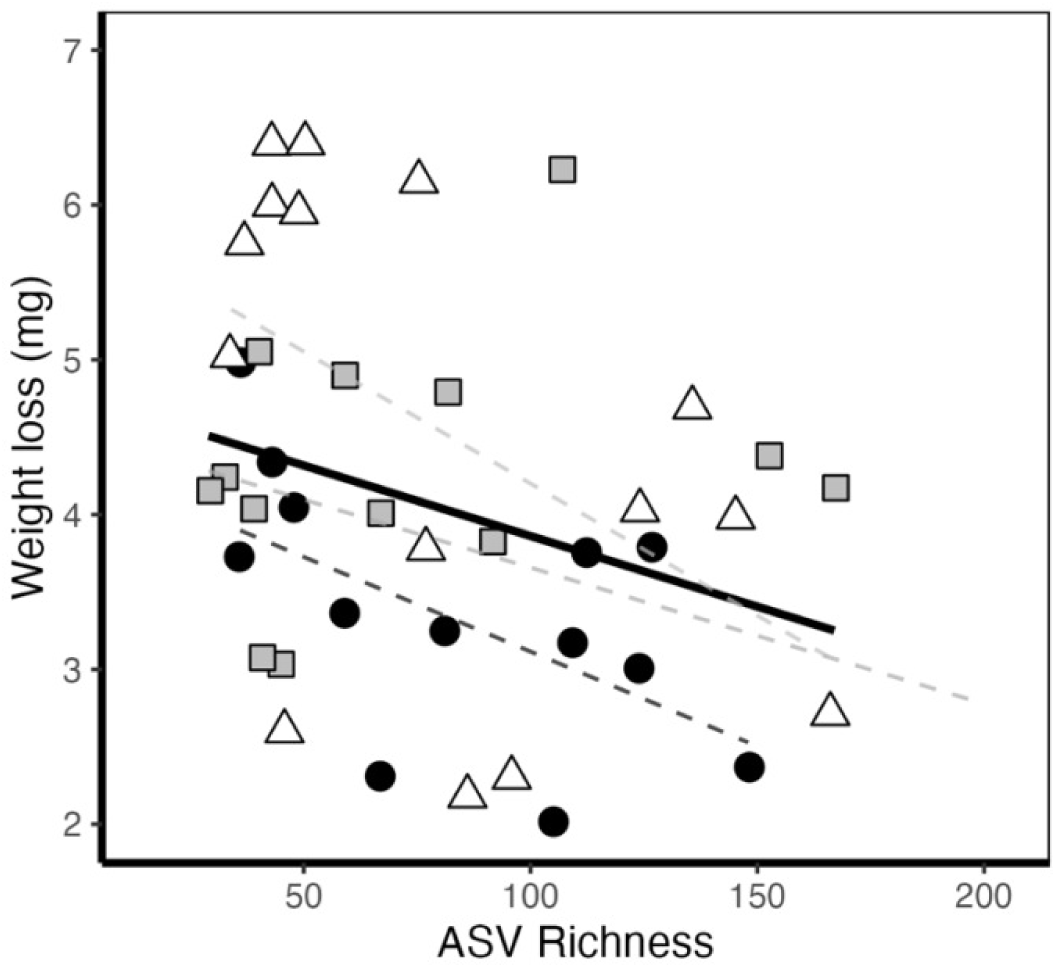
Negative relationship between ASV richness and prey weight loss. Lines indicate best fit lines for the general model (solid black), and independently for each series (dashed). Symbols and line colors represent series 1 (dark grey, black circles), 2 (light grey, grey squares) and 3 (very light grey, white triangles).

**Table 2.** Wald test results for the best performing GLM model, including richness, series, and their interaction.

| Parameter | Sum of Squares | d.f. | F-value | p value |
| --- | --- | --- | --- | --- |
| Richness | 1.244e-05 | 1 | 8.630 | **0.005** |
| Series | 9.785e-05 | 1 | 6.789 | **0.013** |
| Richness : Series | 3.621e-06 | 1 | 0.273 | 0.604 |

Significant p values are indicated in bold.

### Narrow functions

Narrow functions had positive and neutral relationships with diversity. Two functions had significantly positive relationship with diversity: 2-hydroxybenzoic acid and Tween 80 (Figure 2, S3, Table 3). Other carbon sources had positive, albeit weaker, relationships with diversity, and significance was lost after adjusting for multiple testing: including L asparagine, L arginine, D malic acid, and alpha cyclodextrin (Table 3).

**Figure 2.**
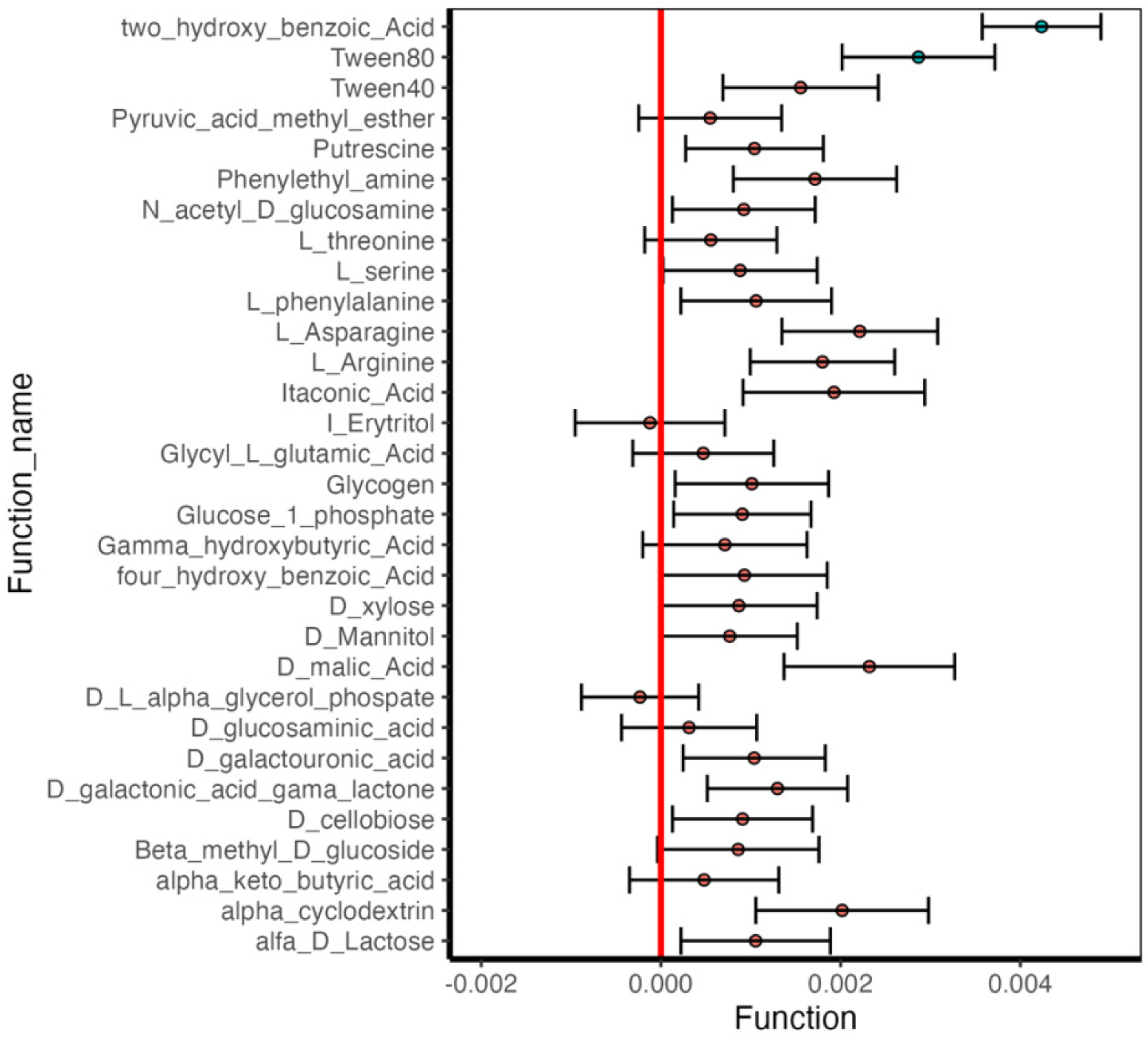
BEF relationship estimates between narrow functions and ASV richness. Values represent estimate of the slope. Fill represents whether the relationship was significant (blue) or not (red) and error bars indicate standard error for each slope estimation.

**Table 3.** GLMM results for diversity correlations with associated narrow functions. Significance is indicated in bold.

| Function name | d.f. | R2 | Estimate | Standard Error | t value | p value | Adjusted p value |
| --- | --- | --- | --- | --- | --- | --- | --- |
| **2 hydroxybenzoic acid** | **3** | **0.489** | **0.004** | **0.001** | **6.412** | **0.000** | **0.000** |
| **Tween80** | **3** | **0.198** | **0.003** | **0.001** | **3.372** | **0.002** | **0.025** |
| Tween40 | 3 | 0.050 | 0.002 | 0.001 | 1.798 | 0.080 | 0.274 |
| Pyruvic acid methyl ester | 2 | 0.000 | 0.001 | 0.001 | 0.689 | 0.494 | 0.589 |
| Putrescine | 2 | 0.000 | 0.001 | 0.001 | 1.360 | 0.181 | 0.434 |
| Phenylethyl amine | 3 | 0.057 | 0.002 | 0.001 | 1.887 | 0.066 | 0.257 |
| N acetyl D glucosamine | 2 | 0.000 | 0.001 | 0.001 | 1.162 | 0.252 | 0.434 |
| L threonine | 2 | 0.000 | 0.001 | 0.001 | 0.755 | 0.455 | 0.564 |
| L serine | 2 | 0.000 | 0.001 | 0.001 | 1.030 | 0.309 | 0.457 |
| L phenylalanine | 2 | 0.000 | 0.001 | 0.001 | 1.264 | 0.213 | 0.434 |
| L Asparagine | 3 | 0.116 | 0.002 | 0.001 | 2.553 | **0.015** | 0.148 |
| L Arginine | 3 | 0.087 | 0.002 | 0.001 | 2.236 | **0.031** | 0.191 |
| Itaconic Acid | 3 | 0.059 | 0.002 | 0.001 | 1.905 | 0.064 | 0.257 |
| I Erytritol | 2 | 0.000 | 0.000 | 0.001 | -0.146 | 0.885 | 0.885 |
| Glycyl L glutamic Acid | 2 | 0.000 | 0.000 | 0.001 | 0.601 | 0.551 | 0.626 |
| Glycogen | 2 | 0.000 | 0.001 | 0.001 | 1.185 | 0.243 | 0.434 |
| Glucose 1 phosphate | 2 | 0.000 | 0.001 | 0.001 | 1.185 | 0.243 | 0.434 |
| Gamma hydroxybutyric Acid | 2 | 0.000 | 0.001 | 0.001 | 0.779 | 0.440 | 0.564 |
| 4 hydroxy benzoic Acid | 2 | 0.000 | 0.001 | 0.001 | 1.010 | 0.319 | 0.457 |
| D xylose | 2 | 0.000 | 0.001 | 0.001 | 0.998 | 0.324 | 0.457 |
| D Mannitol | 2 | 0.000 | 0.001 | 0.001 | 1.020 | 0.314 | 0.457 |
| D malic Acid | 3 | 0.106 | 0.002 | 0.001 | 2.441 | **0.019** | 0.148 |
| D L alpha glycerol phosphate | 2 | 0.000 | 0.000 | 0.001 | -0.356 | 0.724 | 0.748 |
| D glucosaminic acid | 2 | 0.000 | 0.000 | 0.001 | 0.417 | 0.679 | 0.726 |
| D galactouronic acid | 2 | 0.000 | 0.001 | 0.001 | 1.312 | 0.197 | 0.434 |
| D galactonic acid gama lactone | 3 | 0.040 | 0.001 | 0.001 | 1.662 | 0.104 | 0.323 |
| D cellobiose | 2 | 0.000 | 0.001 | 0.001 | 1.165 | 0.251 | 0.434 |
| Beta methyl D glucoside | 2 | 0.000 | 0.001 | 0.001 | 0.957 | 0.344 | 0.464 |
| alpha keto butyric acid | 2 | 0.000 | 0.000 | 0.001 | 0.579 | 0.566 | 0.626 |
| alpha cyclodextrin | 3 | 0.075 | 0.002 | 0.001 | 2.099 | **0.042** | 0.217 |
| alfa D Lactose | 2 | 0.000 | 0.001 | 0.001 | 1.268 | 0.212 | 0.434 |

Narrow functions and the broader function of degradation seemed negatively related (Fig 3, S4), but none of these relationships were significant. D-galacturonic acid and four-hydroxy benzoic acid had the strongest negative relationships with weight loss but significance was lost with corrections (Table S3). In contrast, we found many instances of positive correlations between narrow functions (Figure 3). Notably, 2-hydroxybenzoic acid and Erythritol had fewer correlations than other functions and seemed to associate to different taxa.

**Figure 3.**
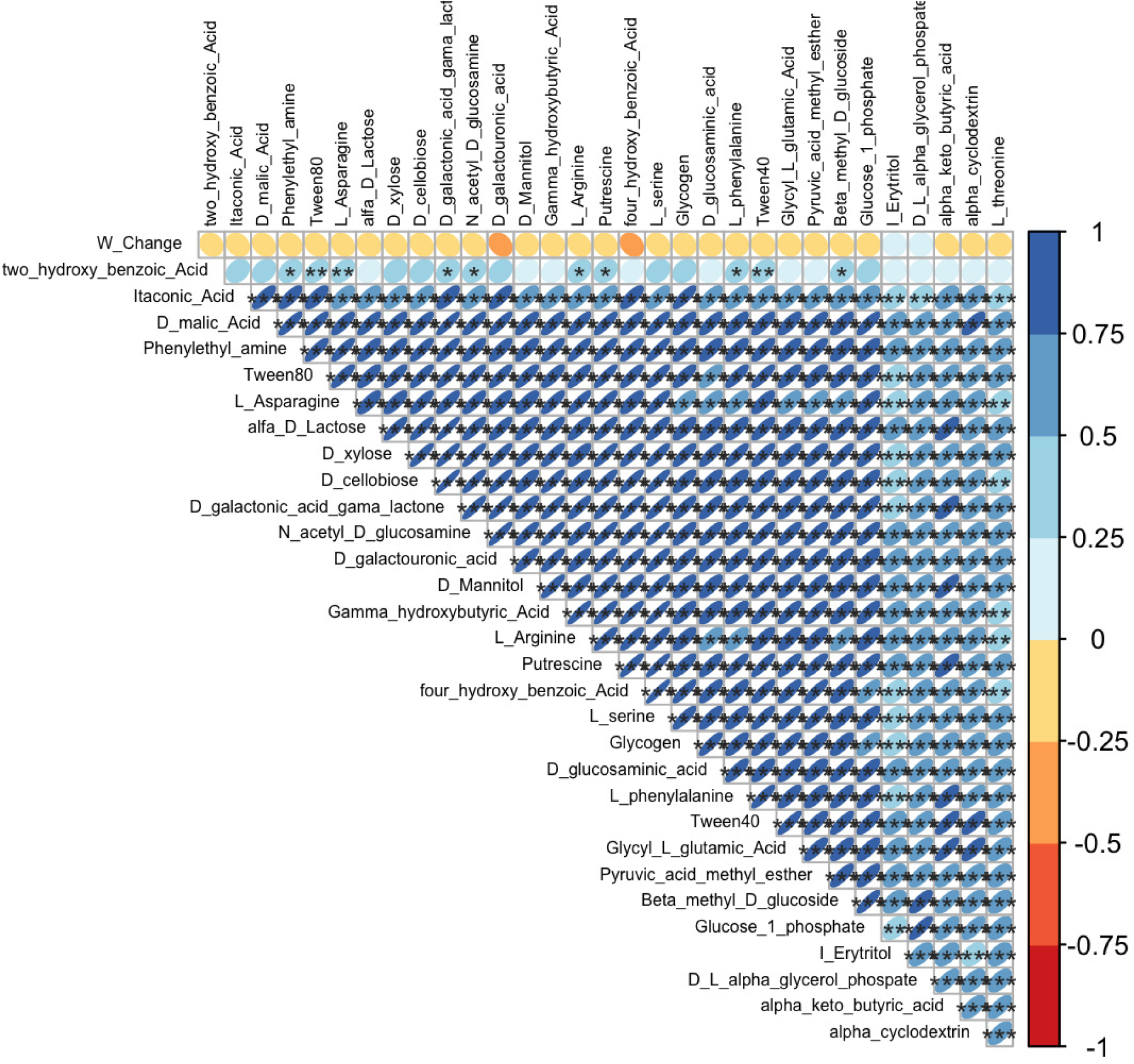
Correlations between functions. Colors and ellipses indicate correlation values while asterisks indicate significance at the 0.05 (*), 0.01 (**), and 0.001 (***) p values.

We identified 3 taxa with significant associations with weight loss (Fig 4, Table S4). *Clostridium* sp. (Clostridiaceae) had a positive association with weight loss while *Dyadobacter* sp. (Spirosomaceae) and *Herbinix* sp. (Lachnospiraceae) had a negative association. *Herbinix* sp. had the strongest correlation with R^2^ of 0.52 (Table S4). Additionally, there were 13 associations between taxa and narrow functions related to carbon flow, all by ASVs within the 100 most abundant (Table S4). Specifically, four taxa associated positively with Tween 80 and eight with alpha cyclodextrin, five positively and three negatively. Despite the significant trends, slopes remained shallow.

**Figure 4.**
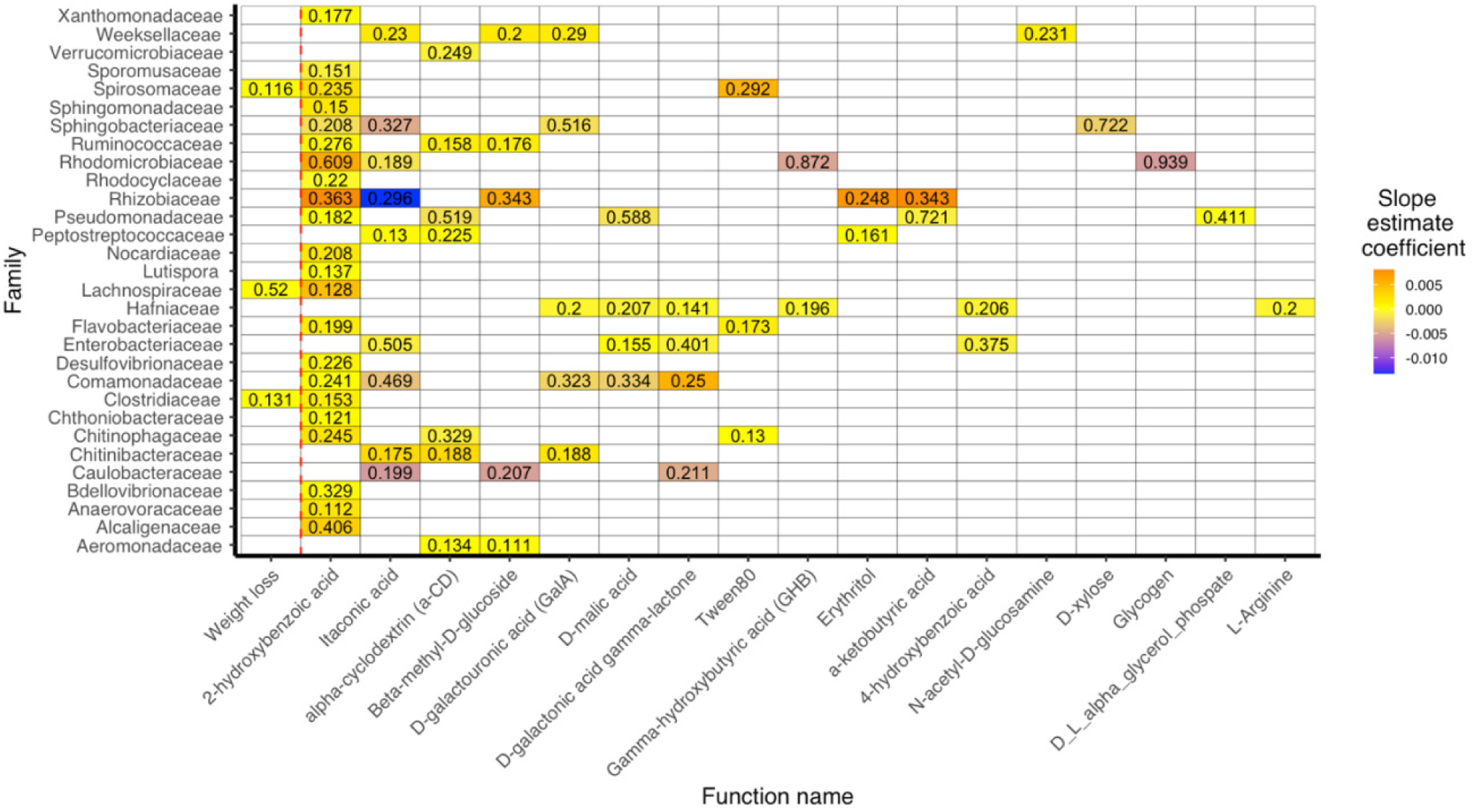
Heatmap showing the correlations between narrow functions and select ASVs. Only significant relationships are shown (adjusted p values < 0.05). Relationships were assessed with a linear model. The fill color indicates the slope estimate and the value within each cell indicates the adjusted R^2^. The red dashed line separates the broad function of degradation measured as weight loss from the narrow functions of carbon utilization.

We found stronger associations between taxa and narrow functions at the level of family (Fig 4). Some substrates are notable due to the large number of associations with different bacterial families, such as two-hydroxy benzoic acid or Itaconic acid. These two substrates also had the strongest associations with a taxon (Rhyzobiaceae), positive for 2-hydroxybenzoic acid and negative for itaconic acid. Five functions were associated with one taxon (right hand side of Fig 4), and these taxa were always able to perform other functions. In contrast, several taxa utilized only one carbon source, namely, 2-hydroxybenzoic acid. In conclusion, we found a diverse array of associations across narrow functions and taxa.

Community composition changed along the dilution gradient and due to series effects (Table 5, Fig 5). The change was progressive and slowed down after the third dilution. The weight change was correlated with the first axis of the NMDS, and richness was not included in the best performing model (Table S5, Pearsons=-0.373, Wald test: F_1_=13.92, p<0.001). We identified 78 taxa significantly associated with diversity gradients after corrections for multiple testing (Table S5) out of which 58 ASV were positively associated with diversity and 20 were negatively associated. Forty-three were also associated with function (Table S4), some positively (31 ASVs), some negatively (12 ASVs). We reviewed the taxa previously associated with function (Table 4) and found that *Dyadobacter* sp (ASV19) had a positive and significant correlation with diversity, whereas *Herbinix* had a slightly weaker correlation (Table S5). *Clostridium* sp (ASV6) had an extremely weak association with diversity. To further establish the role of composition in driving functional rates, we showed that the first axis of the NMDS was associated with weight loss (Table S6).

**Figure 5.**
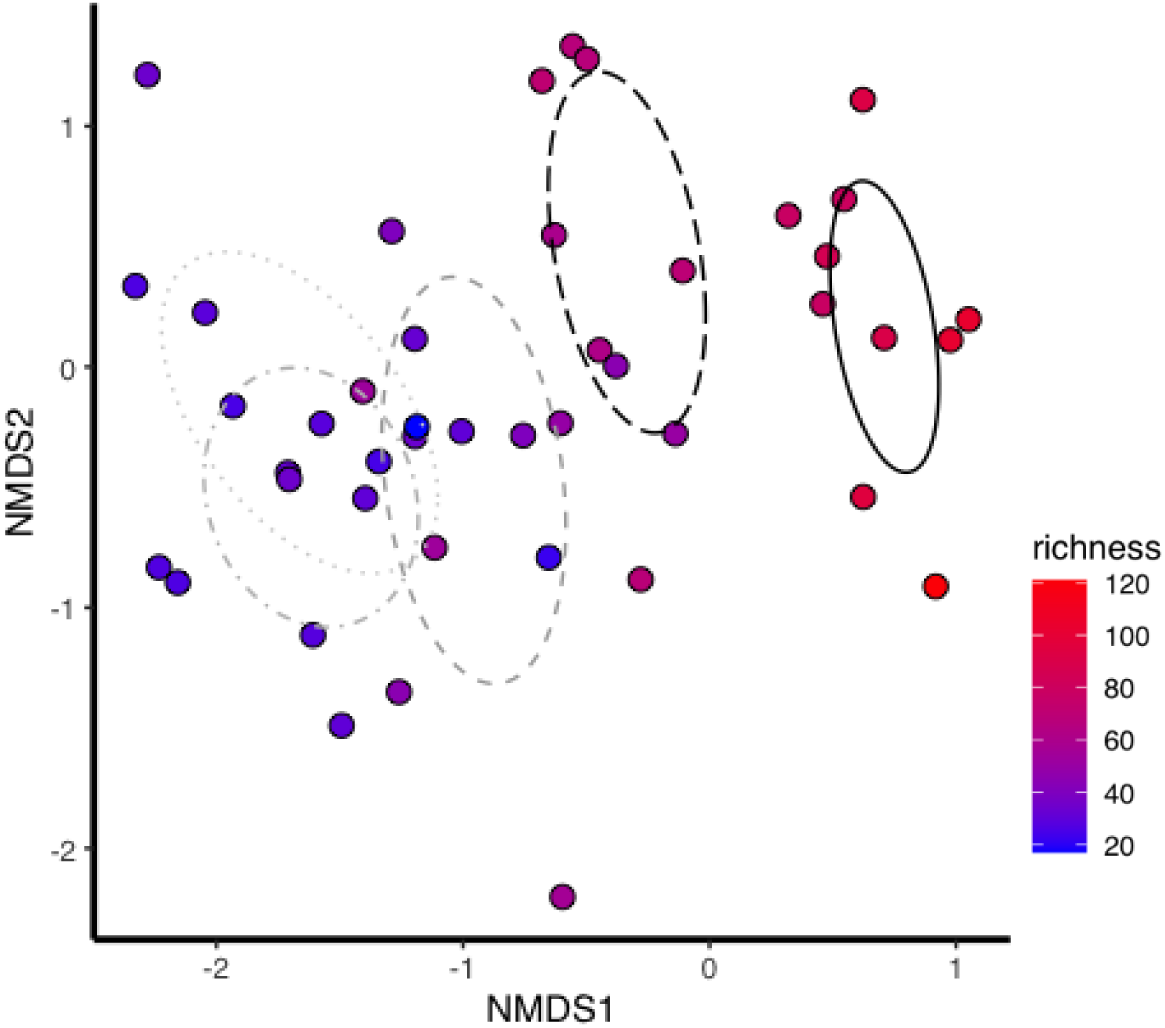
NMDS illustrating the change in community composition. Color reflects richness of each sample. Lines depict the covariance for each dilution group with solid black being the least diluted and dotted light grey the most diluted. Stress was 0.1005.

**Table 4.** GLM model results for ASVs with significant associations with weight loss.

| Taxon |  | df | Estimate | Standard Error | t value | p value | Adjusted p value | R2 |
| --- | --- | --- | --- | --- | --- | --- | --- | --- |
| Clostridium ASV6<br>(Clostridiaceae)<br>(1.23%) | Intercept | 2 | 0.0028 | 0.0002 | 9.663 | <0.001 | <0.001 | 0.130 |
|  | slope | 2 | 9.30E-07 | 2.49E-07 | 3.732 | <0.001 | 0.0284 |  |
| Dyadobacter ASV19<br>(Spirosomaceae)<br>(0.38%) | Intercept | 2 | 0.0043 | 0.00018 | 23.082 | <0.001 | <0.001 | 0.116 |
|  | slope | 2 | -1.33E-06 | 3.79E-07 | -3.509 | <0.001 | 0.0309 |  |
| Herbinix ASV90<br>(Lachnospiraceae)<br>(0.02%) | Intercept | 2 | 0.0051 | 0.0004 | 11.288 | <0.001 | <0.001 | 0.519 |
|  | slope | 2 | -2.03E-05 | 4.60E-06 | -4.406 | <0.001 | 0.0284 |  |

**Table 5.** Adonis. PerMANOVA assessing the effect of dilution factor and ASV richness as an explanatory factors for community composition. Significant p values are indicated in bold.

|  | df | Sum of squares | R2 | F value | p value |
| --- | --- | --- | --- | --- | --- |
| **Series** | 1 | 0.1592 | 0.031 | 1.974 | 0.073 |
| **Dilution factor** | 1 | 1.6112 | 0.315 | 19.977 | **0.001** |
| **interaction** | 1 | 0.1162 | 0.022 | 1.440 | 0.189 |

### Field sites

From pitcher plant fluid collected at the three field sites, we obtained 228,489 reads, with an average of reads/sample 19,040.75. We found 15 bacterial Phyla, accounting for 136 Families and 218 Genera. We lost 4 samples due to low quality sequences or low read numbers, and kept 14 samples for analysis. Field samples had consistently and significantly higher diversity when compared with experimental data (Fig 6, Kruskal-Wallis, χ^2^=46.046, d.f.=2, p<0.001). Field samples averaged 188.17 (± 66.28) ASVs, which was more than twice the mean richness in experimental data (81.31 ± 45.22 ASVs). Field samples were also more even than experiment data (Pielou’s evenness of 0.81 in field and 0.602 in the experiments). Samples taken after the degradation were consistently lower in diversity than any other dataset (ASV richness of 28.05 and Pielou’s evenness of 0.43). Field samples were also significantly different in composition than experimental samples taken before or after the degradation assay (Fig 6c and d, perMANOVA, F=46.207, df=2, R2=0.558, p=0.001). Samples from the field were remarkably similar to one another in composition (Fig 6d, S5). Importantly, representatives of families Clostridiaceae, Spirosomaceae, and Lachnospiraceae were found across most field samples (Fig S6, Lachnospiraceae was found from one of the Plumas sites), indicating the *Clostridium* sp – *Dyadobacter* sp - *Herbinix* sp. relationship we found in experimental samples may be representative of taxa in wild pitcher plants.

**Figure 6.**
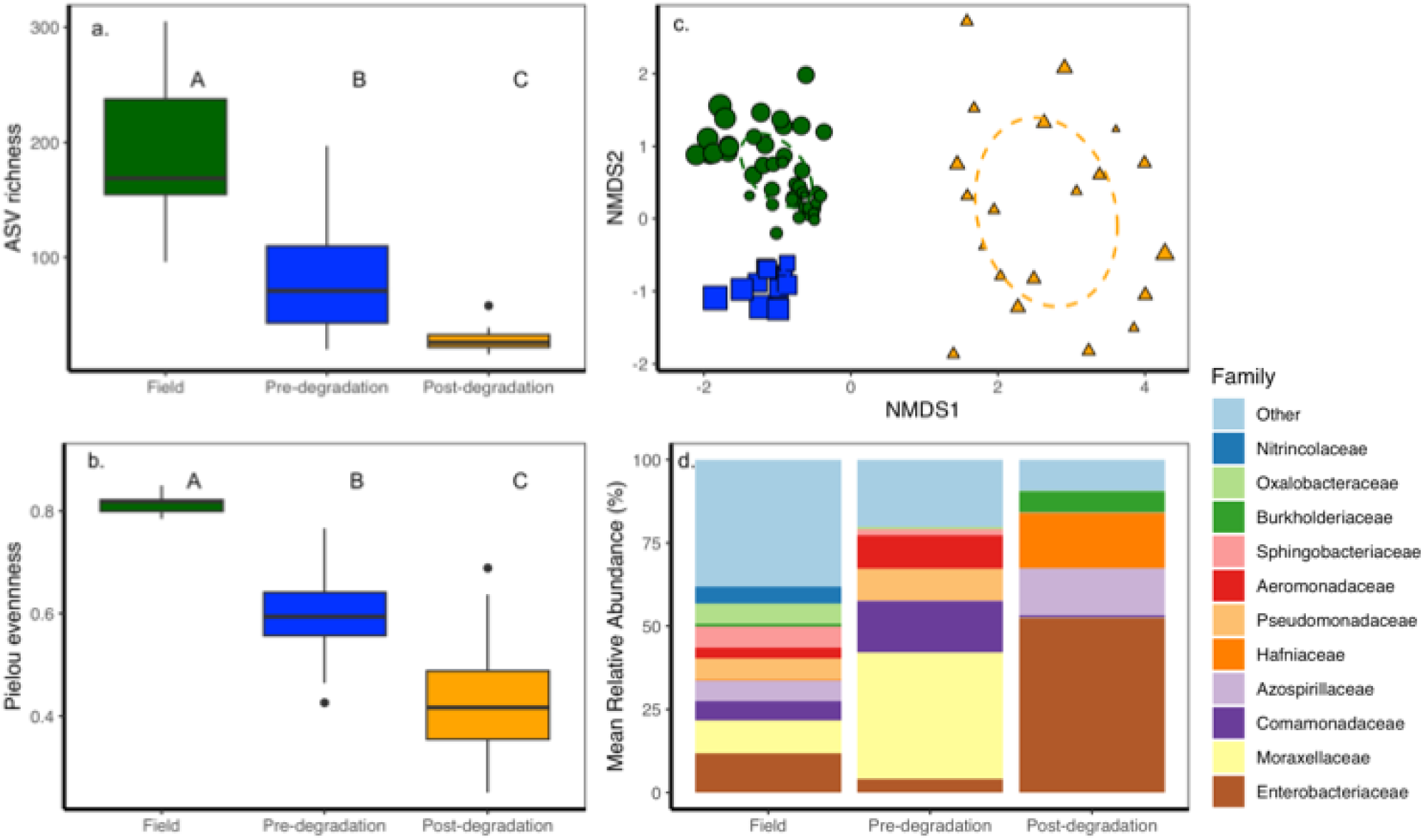
Pitcher plant fluid bacterial communities from field samples. To compare with Experimental data, we calculated richness and evenness (a, b), built an NMDS (c) and a barplot with family level relative abundance (d). Colors (a,b and c) indicate the dataset (Field, and experimental – before and after the degradation assay). Letters (a,b) represent significant differences between groups. Dashed lines (c) correspond to standard error for each dataset. Barplot colors (d) represent family level relative abundance.

## Discussion

Biodiversity – ecosystem function (BEF) relationships are key to connect community assembly to ecosystem function [3, 94]. However, accurately predicting or managing the direction of these relationships remains challenging because we lack a complete understanding of how function emerges from community dynamics[95]. In this study, we classified functions as either broad or narrow based on their scope, to examine whether this categorization offers new insights into BEF dynamics and predictability. Our findings expand the landscape of BEF relationship across functional categories. We found that prey degradation in pitcher plants was negatively associated with bacterial richness, and unrelated to carbon use. This negative BEF relationship is likely driven by subordinate species like *Clostridium* sp. with large functional contributions to decomposition. From species abundance distributions and function-function relationships, we present a preliminary model for prey degradation in pitcher plants that incorporates species individual functional role and community level interactions. We propose this approach could represent a generalized framework to understand the BEF relationships for complex functions.

The negative correlation between bacterial richness and broad functions like degradation contrasts with theoretical expectations and is the first report of negative BEF relationships in the context of broad-narrow functions. There are only three other studies focusing on plant biomass that explicitly show negative BEF relationships, attributing the pattern to slow growing but competitively dominant trees in managed forest stands in Quebec [96], priority effects of biomass accumulating grasslands in Michigan [97], and evidence of negative selection effects in Texas grasses [98]. Additionally, our findings coincide with previous work reporting negative BEF relationships in pitcher plants, including a re-analysis of BEF in experimental bacterial communities [11, 99] and a field experiment showing declining bacterial diversity and increasing degradative function as pitcher plants age [100]. Despite this evidence, negative BEF relationships remain rare in the literature, either because they are fundamentally rare, or because we have not found many examples (van der Plas 2019, D’Andrea et al 2023). To help disentangle these possibilities, we use the data from narrow functions and species relative abundances to establish the underlying mechanisms of negative BEFs, and evaluate assumptions from the literature (Table 1).

Carbon use functions were unrelated to prey degradation, suggesting carbon use and prey weight loss are decoupled. Because all carbon enters via prey [60], this independence suggests limited competition for carbon sources, and that other narrow functions may limit degradation at our scale [12, 101]. Although identifying the rate-limiting pathway is beyond our scope, it likely involves enzymes breaking down integument, cuticle, chitin, fatty acids, or protein into simpler compounds [57, 102]. Once in simpler form, microbial communities can use substrates via complementarity or functional redundancy resulting in efficient and resilient communities [103–106]. Specifically in pitcher plants, declines in bacterial diversity and abundance can influence specific functions such as chitinase activity and prey weight loss [100, 107]. If substrates have different relationships with microbial diversity, accounting for these differences may support our ability to predict how microbial function may change in the face of disturbance [51, 103].

Specifically, we found 2 positive and 29 neutral BEF relationships in the narrow functions we studied (Fig 2). This variation in slopes challenges earlier expectations for narrow functions, where positive relationships were anticipated due to the reliance on specific taxa to perform these specialized functions. Our result coincides with more recent work recognizing neutral BEF relationships in narrow functions (Table 1), and with a broader view on the variety of BEF relationships found in nature [7]. High levels of redundancy coincide with previous studies in soil and aquatic habitats (Table 1). Therefore, narrow functions underlying degradation may be a representative sample of BEF relationships occurring in other systems.

Two narrow functions with positive BEF relationships, 2-hydroxybenzoic acid and Tween80, may influence microbe-plant relationships within pitcher plant leaves. Hydroxy benzoic acids are a polyphenol, common in antibacterial plant metabolites [108, 109]. Antimicrobial compound breakdown in soil and water is well studied for pollution mitigation [110, 111]. Therefore, the many taxa associated with 2-hydroxybenzoic acid utilization (Figure 4) and their diverse enzymes [109] suggest that functional complementarity may underlie its positive BEF relationship with bacterial richness (Fig 4). In contrast, Tween 80 is an emulsifier made of lipid components commonly found in plants [35], and cellulase enzymes from microorganisms can break this down into more bioavailable lipids [112]. Thus, Tween utilization is predicted to be a specialized source of carbon, requiring specific exoenzymes that then make lipids available to the broader microbial community [112, 113]. We found only three Families associated with Tween 80 utilization, and speculate that its positive relationship with diversity may be related to the benefits of lipid availability to taxa unable to utilize this resource directly. The BEF relationships for narrow functions seem thus to relate to specific dynamics within the bacterial community, highlighting the complexity of distinct natural histories in driving patterns that may seem alike at first glance.

We found three taxa that correlated with prey weight loss, identified as *Dyadobacter* sp., *Herbinix* sp. and *Clostridium* sp. *Dyadobacter* sp. and *Herbinix* sp. had negative associations with prey weight loss, suggesting they have an antagonistic relationship with the broader function. *Herbinix* sp. stood out because of its strong positive association with diversity. Related organisms have been recorded to participate in degradation in the context of eutrophic soils [114], hydrogen reactors, [115], biogas plants, and wastewater [116]. Yet, *Herbinix* sp. in our dataset was negatively correlated with degradation and with *Clostridium* sp.’s abundance, suggesting an antagonism, such as competition, may be key to understanding the negative BEF relationship of degradation. Although these specific genera were not detected in field samples, members of the same families (Clostridiaceae, Spirosomaceae, and Lachnospiraceae) were consistently found across field sites (Fig 6, S5, S6), suggesting these dynamics may be general to these pitcher plant microbial communities.

In addition to its positive association with prey weight loss, this *Clostridium* sp. had relatively low abundance in the microcosms, and was negatively associated with diversity. This taxon is thus a strong candidate to underly the negative BEF in the broad function. Because its association with narrow functions is indistinguishable from other taxa profiles, this *Clostridium* sp. likely performs another specific function, key and rate limiting for insect degradation. Closely related taxa perform degradation in lakes[117], wetlands [118], wastewater [119, 120], and earth worm guts [121] with key roles in nitrogen cycling across all these habitats. Broadly, *Clostridium* holds 120 species with a variety of physiological traits related to degradation, strong sporulating ability, and dramatic competitive interactions with related taxa [122, 123]. We speculate that the negative relationship between biodiversity and prey decomposition is explained by one or more degradative functions performed by this *Clostridium* sp. as it is released from competition from related taxa such as *Herbinix* sp.

Although our results have clear implications for our understanding of BEF relationships, there are two limitations that are important to consider. First, we performed measurements at the beginning and end of the degradation assay due to methodological constraints, though these processes may occur in different time scales [124–126]. Taxonomic and functional shift during animal decay often follow predictable steps, linked to substrate changes, as observed in human corpses [124]. Based on our robust findings, we argue that a snapshot of the time-series of degradation is sufficient to establish BEF relationships for narrow and broad functions. Second, relying on correlations to link function to function and functions to community structure prevents us from establish causative relationships. Others have used post-hoc experimentation [15], and metagenomics [127, 128], to further link community structure and functional outputs.

In this study, bacterial diversity and degradation had a negative relationship because key functional contributors to degradation are subordinate species that perform better in low-diversity. In wild pitcher plants, this relationship suggests that leaves with low bacterial diversity may degrade prey faster than those with high bacterial diversity. Because a single plant can hold many leaves, pitcher plants are likely able to deal with varying microbial activity by mobilizing nitrogen from productive leaves to growing tissues [129]. This variation in microbial community diversity and composition can be driven by leaf age [57, 100], historical contingencies [99], prey capture patterns [130], or across sites if we consider biogeographical variation (Fig 6).

Our findings highlight the complexity of establishing BEF relationships across functions. Importantly, we found negative BEF in broad functions and non-negative shapes in narrow functions, challenging the traditional expectations across these categories (Table 1). Thus, our study suggests the landscape of BEF relationships across narrow and broad functions is diverse. Additionally, accounting for function-function relationships and species contributions helped establish a scenario for subordinate species functional contributions and discard simple carbon substrate utilization as drivers of degradation or competitive interactions. Understanding the relationship between coexistence mechanisms and complex functions like degradation should further advance our ability to predict ecosystem change [103, 131].

## Data availability statement

Raw sequences were deposited in the NCBI Sequence Read Archive and can be found under accession number PRJNA1287784. All code and metadata are available in the Github page https://github.com/catalicu/DegExp2019.

## Acknowledgements

We thank Brianne Lee for facilitating access to the Greenhouse. We thank Susan Wright for laboratory support. We thank Six Rivers National Forest, Plumas National Forest, and Shasta Trinity National Forest for permits, support and access to pitcher plant populations. Support for this project was obtained by NSF DEB 2046214 awarded to CCG.

